# Heatwaves do not impact bacteria within pollen provisions, despite accelerating blue orchard bee (*Osmia lignaria*) larval development

**DOI:** 10.64898/2026.08.26.747351

**Authors:** Alexia N. Martin, Neal M. Williams, Rachel L. Vannette

## Abstract

Many insect populations are experiencing thermal stress as a result of global change, making it imperative to investigate how their relationship with other organisms will be impacted by heat disturbances. Microbial symbionts, such as bacteria, have the potential to enhance or inhibit an insect’s thermal tolerance. Solitary bee larvae host bacteria within their food stores (“pollen provisions”), which have been shown to benefit survival and development; however, it is unclear how heatwaves brought about by climate change will impact their relationships with these bacterial partners. In this study, we subjected blue orchard bee (*Osmia lignaria*) eggs and larvae to a 4-day heatwave (35 °C daytime:22 °C nighttime) or kept them at control temperatures (25 °C daytime:15 °C nighttime), then returned all bees to control temperatures for a 5-day recovery period. We assessed bacterial communities within pollen provisions and larval development stage pre-heatwave (Day 0), immediately post-heatwave (Day 4), and following the recovery period (Day 9). Bacterial community composition, diversity, and abundance were resilient to heat stress, but larval bees developed faster when subjected to a heatwave. This finding refutes the hypothesis that bacteria within pollen provisions modulate blue orchard bee responses to heat, suggesting instead that developmental effects could be more largely shaped by bee physiology or interactions with microorganisms other than bacteria.

## Introduction

Given rapidly warming temperatures and altered precipitation regimes, it is critical to understand how changing climate conditions will shape future interactions between insects and other organisms. Although insects are one of the most diverse taxa on Earth with an estimated 14 to 20 million species (Colwell et al. 2026), ongoing climate change likely threatens hundreds of thousands of species (Cardoso et al. 2020), including those that provide critical ecosystem services like native bees (Potts et al. 2010, Soroye et al. 2020, LeBuhn and Vargas Luna 2021). Extreme weather events, such as heatwaves, are increasing in intensity and frequency, exposing bees to unprecedented stress (Meehl et al. 2007, Ummenhofer and Meehl 2017). These extreme temperatures can push individual bees outside of their optimal thermal range, resulting in death, and ultimately lead to population declines, range shifts, and changes in community composition (Kuhlmann et al. 2012, CaraDonna et al. 2018, Soroye et al. 2020, Kazenel et al. 2024, Melone et al. 2024, Vilchez-Russell and Rafferty 2024).

Symbiotic interactions between insects and microorganisms may enhance or inhibit an insect’s ability to survive in stressful environments (reviewed by Lemoine et al. 2020). For instance, microbial partners can expand insect diet-breadth (Cornwallis et al. 2023), increase desiccation resistance (Engl et al. 2018), and buffer against heavy metal exposure (Rothman, Leger, et al. 2019). Thermal tolerance may also be linked to their microbial partnerships (reviewed by Renoz et al. 2019), which has particular relevance for climate resilience. Insects that host obligate mutualists may have a microbial Achilles’ heel (e.g., Kikuchi et al. 2016, Renoz et al. 2019), as heat stress can reduce symbiont abundance and result in decreased host fitness (Dunbar et al. 2007, Burke et al. 2010, Kikuchi et al. 2016, Shan et al. 2017); however, facultative symbionts may be less sensitive to heat stress (Shan et al. 2017) and in some cases can buffer their hosts to negative heat effects (Burke et al. 2010, Gruntenko et al. 2017). Therefore, characterizing extreme heat impacts to bees’ microbial symbionts must also be considered to fully understand how bees will be affected by these aspects of climate change.

For above-ground nesting solitary bees, eggs and larvae may be especially vulnerable to heatwaves because they are minimally buffered against outside temperatures as in social bee species and ground nesting bees, and they are unable to move to new locations to regulate their body temperature. For instance, simulated heatwaves that mimic realistic future California temperatures significantly increased the mortality of blue orchard bee (*Osmia lignaria*) larvae (Melone et al. 2024). Additionally, sustained warming of blueberry bee (*Osmia ribifloris*) larvae during development increased mortality rates, reduced fat content, and delayed adult emergence (CaraDonna et al. 2018). The effects of heatwaves during bee development can even carry over to adulthood, with *Osmia bicornis* exposed to extreme heat as larvae having reduced body mass and impaired flight performance as adults (Gudowska et al. 2026). Although we have some understanding of how these bees will be impacted directly, we are currently missing information as to how realistic heatwave temperatures impact bee-symbiont interactions.

When constructing a nest, solitary bee females give each offspring a mass of pollen and nectar called a pollen provision, which also contains microorganisms like bacteria and fungi. For most solitary bees studied to date, the pollen provision microbiome is reflective of microbes in the environment, containing facultative and commensal partners (Rothman, Andrikopoulos, et al. 2019, Voulgari-Kokota et al. 2020, Nguyen and Rehan 2022, Kueneman et al. 2023, Vannette et al. 2025). These provision-associated microbes can directly confer health benefits to their developing hosts. For instance, larval blue orchard bees and blueberry bees show reduced survival, slowed development, decreased weight, and altered nutrition when reared on sterilized diets or provisions with microbes sourced from different bee species (Dharampal et al. 2019, 2020, 2022). Microbes within pollen provisions have also been suggested to protect larvae against pathogens, such as the case in the bumblebee mimicking digger bee *Anthophora bomboides* (Christensen et al. 2024). Interestingly, experimental attempts to investigate the benefits of single microbial species to megachilid bees through addition to pollen provisions have so far been unsuccessful at uncovering positive health impacts (Brar et al. 2024, Martin et al. 2026), which may suggest that whole communities are imperative for successful development or that benefits are context- or species-specific. Whether provision-borne microbes mediate bee response to elevated temperatures remains unclear; therefore, it is critical to better understand how these microbial symbionts may be impacted by heatwaves, as decreases in microbial diversity or abundance could have negative health implications for developing offspring.

Like with macroorganisms, the community composition and abundance of microorganisms can shift due to environmental disturbances (reviewed by Shade et al. 2012). To make predictions as to how bacterial communities within pollen provisions may respond to heat stress, we may look at how microbes within potential source pools (e.g., flowers and nesting materials; Keller et al. 2013, Rothman, Andrikopoulos, et al. 2019, Voulgari-Kokota et al. 2019, Vannette et al. 2025, Martin et al. 2026) respond to adverse conditions. In flowers, bacteria that reside in nectar can be susceptible to thermal stress, as documented by community composition shifts and reduced microbial diversity (Russell and McFrederick 2022, Cecala et al. 2025). Microbial communities in soil, the material used by blue orchard bees to construct nest walls, display divergent responses to heat. Although there is some evidence of soil microbial resilience to heat stress (Boyle et al. 2024), most studies to date find evidence of disturbance resulting in community composition and alpha diversity shifts following heat treatments (e.g., Bérard et al. 2011, Bei et al. 2023) (reviewed in Rocca et al. 2019). Together, this suggests that microbial communities within pollen provisions may be sensitive to disturbances.

In this experiment, we investigated the impact of a short-term heatwave on the community composition and abundance of bacterial symbionts within blue orchard bee provisions and on larval development. We sampled pollen provisions and monitored larval development at the beginning of the experiment, immediately following a four-day heatwave, and after a five-day recovery period at control temperatures. We predicted that heatwave exposure would shift bacterial community composition and abundance, resulting in alternative states or dysbiosis immediately following the disturbance pulse. If bacterial communities were robust to stress, we expected to see (1) no shift in communities (“reistance”) or (2) reversion to the initial microbiome stable state following return to control temperatures (“resiliance”). However, if bacterial communities were sensitive to heat stress, we expected to see a permanent disturbance in the pollen provision microbiome, marked by a difference in microbial communities even after return to non-stressful temperatures.

## Methods

### Study System

*Osmia lignaria* (Say), the blue orchard bee, is a cavity-nesting solitary bee native to North America. Adult females begin nesting in the spring, with each individual creating her own nest. Nests are formed in cavities of a suitable diameter and mud is used to form walls between each individual brood cell. These bees are mass provisioners, meaning that mothers create a large ball of pollen and nectar (called a pollen provision) for the offspring to consume throughout its development. To create the nest, the mother bee starts by making a wall of mud, then creates a pollen provision, lays an egg upon the pollen provision, then makes another wall of mud (Torchio 1989). She will do this throughout the entire length of the nest, usually forming a vestibule (i.e. empty cell) at the front of the nest.

### Experimental Set-Up

We seeded two 6.3 m x 11 m x 2.04 m screen hoop houses at the University of California, Davis Harry H. Laidlaw Jr. Honey Bee Research Facility with *Phacelia tanacetifolia* in November 2024. At full bloom in April 2025, we released adult male and female *O. lignaria* into the hoop houses to begin nesting. We provided bees in each hoop house with a consistent mud source and three wooden nesting blocks lined with removable paper straws. Nesting blocks were housed in a large black plastic bin to protect them from rain and direct sun exposure. We checked blocks daily to monitor for nesting progress and completion from May 3, 2025 to May 14, 2025. We removed completed nests (n = 22) from the hoop houses, marked them with the hoop house of their origin and nest collection date, then brought them to the lab for processing. We recorded environmental temperatures using an Elitech data logger from May 8, 2025 to May 14, 2025, with temperatures ranging from 7.5°C to 41.9 °C outside of the nesting block and 8.7 °C to 39.8 °C within the nesting block.

### Treatments

To investigate how heatwaves impact bacteria within the pollen provision, we exposed all provisions and their corresponding offspring (hereafter called a “brood cell”) to either a 4-day heatwave (35 °C daytime:22 °C nighttime) or kept them at control temperatures (25 °C daytime:15 °C nighttime) for 4 days (Fig. 1). Following this period, we returned the brood cells from both temperature treatments to control temperatures (25 °C daytime:15 °C nighttime) for a 5-day recovery period. Temperatures were set to 14:10 day:night cycles. The heatwave temperatures were modified from Melone et al. 2024 to promote higher bee survival and were based on California central valley expected temperatures as a result of climate change. We also sampled the initial microbiome within a subset of pollen provisions (Fig. 1, Day 0), in order to obtain a bacterial community baseline.

**Figure 1:**
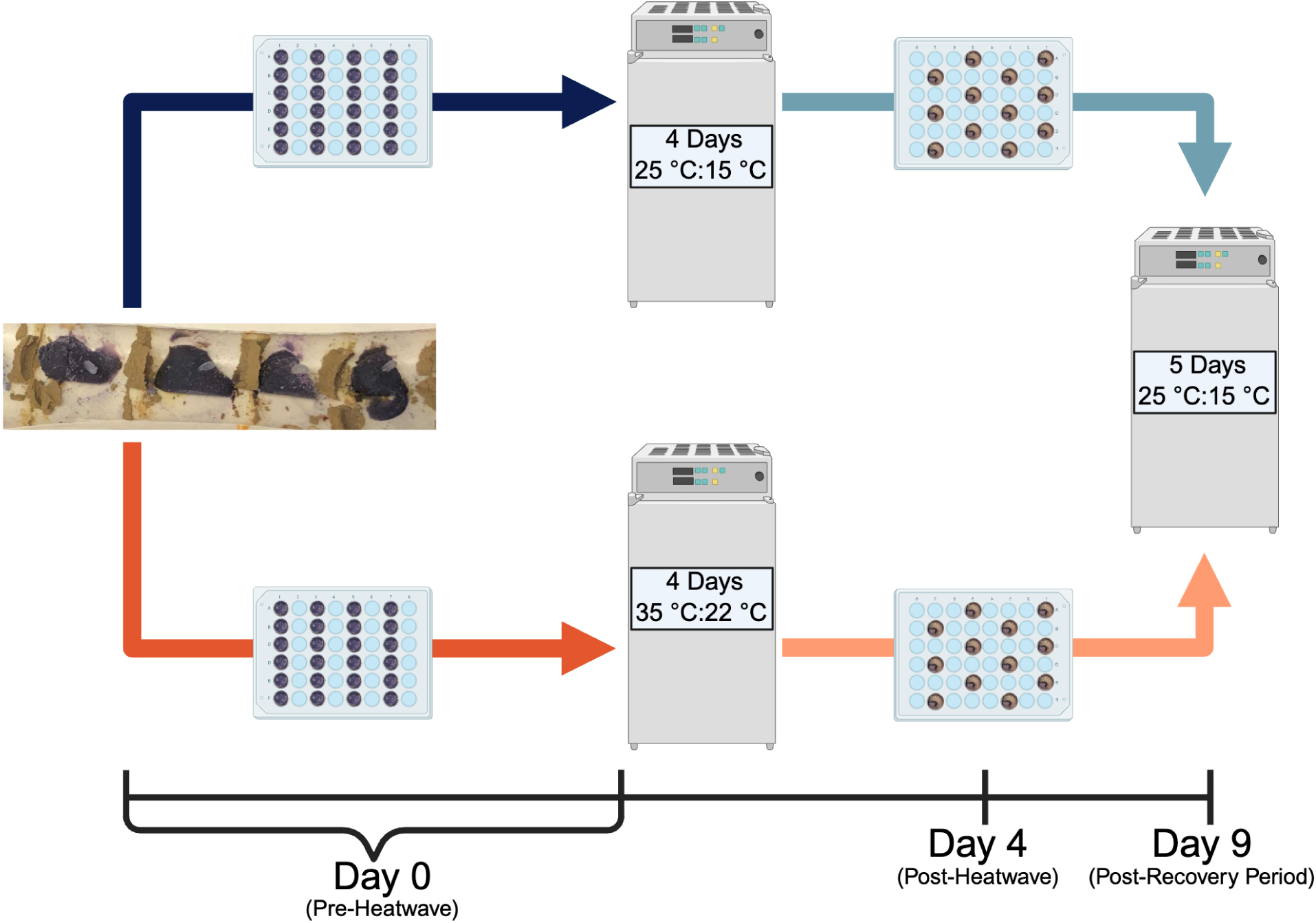
Graphical representation of the study design. Pollen provisions were sampled and larval development was assessed on Day 0 (pre-heatwave), Day 4 (post-heatwave), and Day 9 (post-recovery period). Brood cell photos taken by first-author and figure created in BioRender. <u>Alt Text:</u> Graphical depiction of the experimental design with a timeline to indicate when pollen provision samples were taken and larval development was measured.

### Experimental Application and Sampling

We removed all brood cells (n = 98) from the nesting straw within five days of nest collection, at which point we noted the number of brood cells in a nest, presumed offspring sex (based on position within the nest and size of provision), and initial developmental stage of each offspring. Next, we randomly assigned each brood cell to a treatment (N_Control_ = 38, N_Hot_ = 38), and then placed them into their respective incubators within treatment specific 48-well plates or froze them at −20 °C to sample the pre-heatwave microbiome (N_Pre-heatwave_ = 15). On day 4, we noted the developmental stage of each offspring and froze half of the brood cells in each treatment (N_Control Day 4_ = 20, N_Hot Day 4_ = 19) at −20 °C to sample the pollen provision microbiome immediately post-heatwave. We then moved all remaining brood cells to a control temperature incubator for an additional 5 days, after which we noted the developmental stage of each offspring and froze them at −20 °C to sample the pollen provision microbiome following a recovery period (N_Control Day 9_ = 18, N_Hot Day 9_ = 19). We classified developmental stages as follows: “E” for eggs, “EF” for offspring that were between egg or first-instar, “F” for first instar, “B” for offspring between second and fourth instar, and “FT” for fifth instar. At each time point, we also noted offspring mortality.

### DNA Extraction, Sequencing, and Sequence Processing

We chose a subset of pollen provisions to investigate bacterial diversity in each temperature treatment on all three sampling days (n = 65). To prepare samples for DNA extractions, we weighed ∼1/3 of each pollen provision and added it to a bead beating tube. Next, we extracted DNA from the prepared pollen samples using the QIAGEN DNEasy PowerSoil Pro Kit following kit instructions with one methodological difference – samples were bead beaten using a Benchmark BeadBlaster 24 at speed 7 for four 20 second cycles. We sent the extracted DNA to the Integrated Microbiome Resource located at Dalhousie University in Halifax, NS where the 16S rRNA (V5/V6) region was sequenced using 799F (5′-AACMGGATTAGATACCCKG-3′) and 1115R (5′-AGGGTTGCGCTCGTTG-3′) primers (Chelius and Triplett 2001, Anguita-Maeso et al. 2022, Christensen et al. 2024).

We performed sequence processing and all subsequent analyses in R (version 4.4.1, ‘Race for Your Life’). Using the dada2 pipeline (Callahan et al. 2016), we removed primers from sequences, merged the forward and reverse reads, removed chimeras, and assigned taxonomy using the Silva database (Quast et al. 2013, Callahan 2024). Following initial processing, we used the Phyloseq package to remove mitochondrial (n = 398 ASVs) and chloroplast (n = 1 ASV) reads (McMurdie and Holmes 2013). Next, we confirmed the top 30 most abundant ASVs using the refseq_rna database within NCBI BLAST and a 98% cut-off (Altschul et al. 1990), and noted the isolation source of each ASV’s closest relatives (Table S1). If any ASVs did not have a match present in over 98% identity, we assessed the top five closest matches and assigned the ASV as the lowest level of taxonomic convergence. We used the Decontam package (Davis et al. 2018) to assess contamination based on the prevalence of contaminant sequences across two sequenced kit blanks and the experimental samples (threshold = 0.1). As a result, six ASVs were determined to be contaminants and we removed them from the dataset. Finally, we performed prevalence filtering using the filter_taxa function within the genefilter package, where we set the kOverA value to 2, 2 (Gentleman et al. 2024). Following the dada2 pipeline, samples had an average of 255,596 reads and 94.65% of reads remaining (Table S2). After removal of mitochondria, chloroplasts, and the six contaminant reads, samples had an average of 9,625 reads and 3.6% of reads remaining, with most reads lost after removal of mitochondrial sequences. After prevalence filtering, samples had an average of 9,001 reads and 3.4% of reads remaining. We created sampling curves using the “rarecurve” function in the vegan package (Oksanen et al. 2024) to ensure adequate sampling depth (Fig. S1). Though many reads were lost in processing, the sampling curves fully saturated and the conservative cut-off of 600 reads we chose resulted in all samples being retained. At the end of processing, we created a Phyloseq (McMurdie and Holmes 2013) object that included an OTU table and taxonomy table to use in downstream analysis.

### qPCR

We quantified bacterial rRNA copy number from all samples using qPCR following the same procedure as described in Martin et al. 2026 and detailed briefly as follows. After performing qPCR on a dilution series (1:10 to 1:1000) of a representative subset of samples, we diluted all samples 1:1000 as it yielded Cq values best within the ideal range of 20-30 (25 being optimal). To calculate copy number based on the Cq value, we created a standard curve using a plasmid with the partial 16S rRNA of *Apilactobacillus micheneri* (LC318485.1). We randomized the order of all samples and kit blanks using the R sample() function in order to account for plate effects, with all run in triplicate. Each reaction contained 5 μl of SsoAdvanced Universal SYBR Green Supermix, 3.4 μl of PCR water, 0.3 μl of forward primer 799F (10 μM), 0.3 μl of the reverse primer 1115R (10 μM), and 1 uL of the diluted sample. Thermocycler conditions were set to 95 °C for 3 min, followed by 35 cycles of the following: 30 s at 95 °C for denaturation, 30 s at 52 °C for annealing (fluorophore measured at this step), and 1 min at 72 °C for extension.

### Statistical Analysis

#### Pollen Provision Bacterial Community Analysis

To investigate if overall bacterial communities are impacted by heatwaves, we ran a PERMANOVA using the adonis2() function in R (“vegan” package; Oksanen et al. 2024). In the model, we included nest collection date and the interaction between temperature treatment (Control, Hot) and sampling day (Day 4, Day 9) as predictors and a Bray-Curtis distance matrix as the response variable. For each predictor, we used the betadisper() function to test homogeneity of dispersion. To visualize the bacterial community composition between treatments, we created a Non-metric Multidimensional Scaling (NMDS) plot using a Bray-Curtis Distance matrix. To lower the stress score, we ran the ordination in 3 dimensions. For this and all subsequent analyses, we also tested if the response variable differed over time across our five treatment-sampling categories (Pre-heatwave, Control Day 4, Control Day 9, Hot Day 4, Hot Day 9; see supplemental methods and results). To test if alpha diversity was impacted by heatwaves, we performed a linear model using the lm() function (“stats” package; R Core Team 2024) with the same predictors as described above and Shannon diversity as the response variable. Finally, we performed four separate DESeq analyses to determine if any bacterial genera were differentially abundant across (1) our five treatment-sampling categories, (2) nest collection dates, (3) temperature treatments, or (4) provision sampling days (Love et al. 2014).

#### Pollen Provision Bacterial Abundance Analysis

For each sample, we converted the average Cq value into log copy number using the linear equation from the prepared standard curve. To calculate log copy number per mg of pollen provision, we started by converting the log copy number to copy number by exponentiation. Next, we multiplied the copy number by 50 to account for the total volume of elution buffer the DNA was extracted into and then divided by the mass of pollen provision used for the DNA extraction in milligrams. Because the same primer set was used for DNA sequencing and qPCR, we multiplied this value by the percentage of reads remaining in the sample following prevalence filtering to account for the number of relevant reads remaining in each sample. Finally, we took the log of this value. To determine if heatwaves shift the abundance of bacterial communities in pollen provisions, we ran a linear mixed model with log copy number per mg of pollen provision as the response variable, the same predictors previously described as main effects, and qPCR plate ID as a random effect.

#### Larval Development and Survival Analysis

To investigate if temperature alters development rate, we performed chi-square tests comparing the number of bees at each developmental stage (Egg, Egg/First Instar, First Instar, Between Second to Fourth Instar, Fifth Instar) across the temperature treatments (Control, Hot) using the full dataset. We independently conducted tests for each day development was measured (Day 0, Day 4, Day 9), resulting in three chi-square tests. To investigate if survival was impacted by heat stress, we performed a general linear model using the glm function (“stats” package; R Core Team 2024) with temperature treatment and sacrifice day as predictors and survival status (dead, alive) as the response variable.

## Results

### Bacterial communities in pollen provision were shaped by nest collection date, not heatwave

Bacterial communities were variable across all treatments, with the top 30 most abundant taxa only comprising on average 26% of the relative proportional abundance per sample (Fig. 2). Of these taxa, 27/30 had closest matches isolated from non-bee and non-flower sources (e.g., soil, animal feces, plant roots, and water; Table S1), while the other 3/30 had closest matches with no isolation source listed. Additionally, bacterial community composition (Table 1) and alpha diversity did not differ significantly across temperatures (F = 1.03, df = 1, p = 0.32), sampling days (F = 0.01, df = 1, p = 0.92), or the interaction between the two (F = 1.34, df = 1, p =0.24) (Fig. 3). Instead, nest collection date significantly affected community composition and explained approximately 16% of community variation (Table 1), though variance among dates differed (betadisper F = 7.52, df = 4, P = 0.001). Nest collection date also significantly impacted alpha diversity (F = 3.09, df = 4, p = 0.02). There were nine differentially abundant taxa across the five temperature treatment-sampling categories, eight of which increased in relative abundance from Day 0 (pre-heatwave) to Day 9 (post-recovery period) and one which decreased in abundance over time (Fig. S2); however, there were no taxa identified as differentially abundant across nest collection dates, temperatures, or provision sampling days.

**Figure 2:**
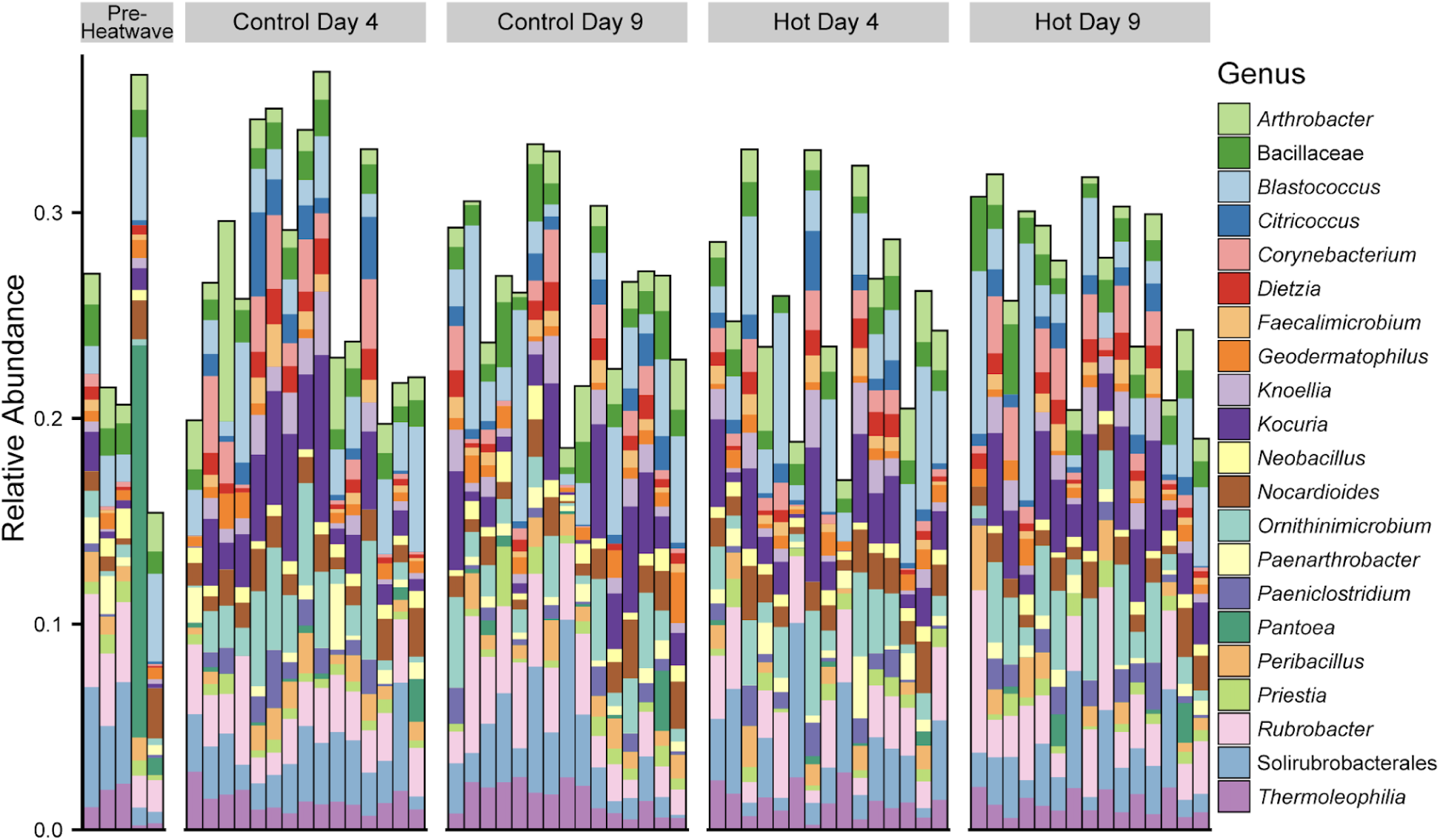
The relative proportional abundance of the top 30 most abundant ASVs within samples at all three sampling points, separated by temperature treatment. ASVs are colored by genus or the lowest level of taxonomic convergence when BLASTed. <u>Alt Text:</u> Stacked bar plot showing the relative proportional abundance of the top 30 most abundant ASVs within each sample. Samples are organized based on their temperature treatment (Control or Heatwave) and when they were sampled (Day 0, Day 4, or Day 9). Across all treatments and sampling periods, stacked bar plots show very similar trends.

**Figure 3:**
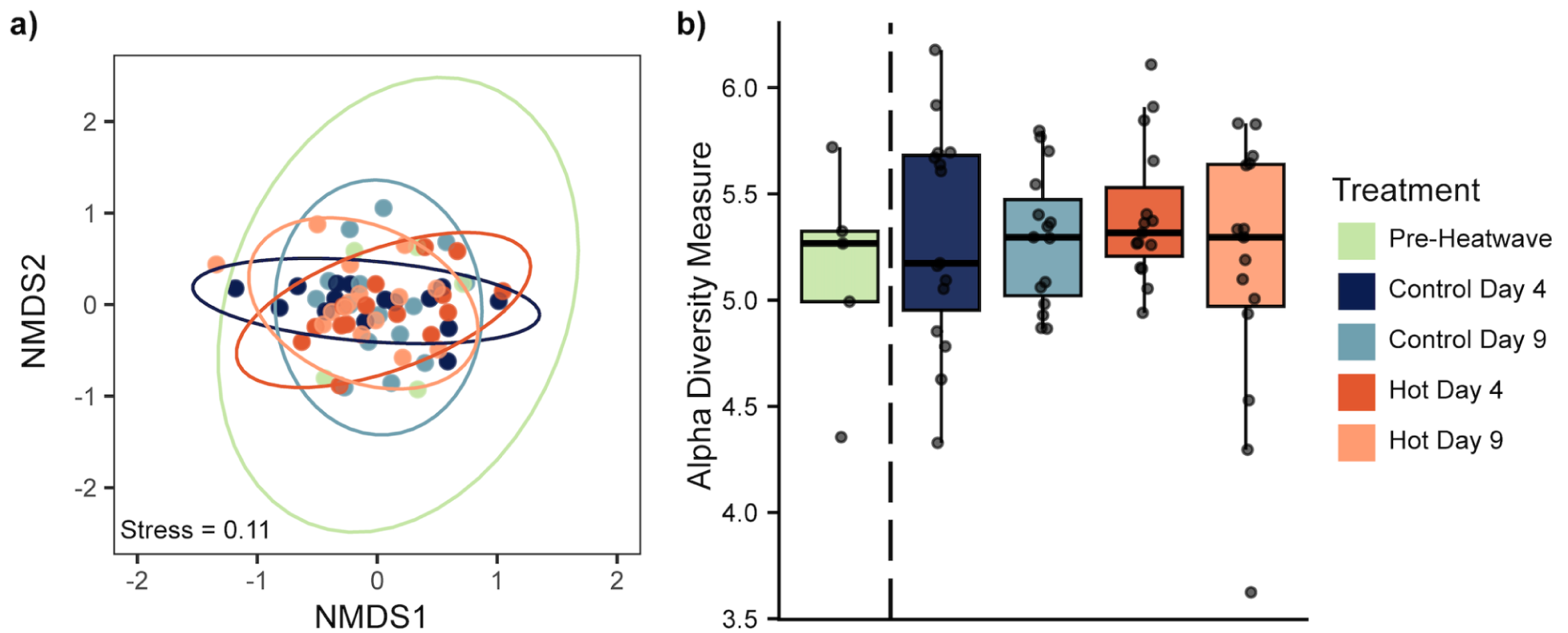
(a) NMDS plot based on Bray-Curtis Distance Matrix and (b) Alpha diversity plot based on Shannon Diversity, colored by temperature treatment and sampling day. Pre-heatwave samples (light green) are included in the plots for visual representation of the initial bacterial community, but we did not include these in the analysis. <u>Alt Text:</u> Two graphs labeled A and B. Graph A is an NMDS ordination with all samples depicted as dots, colored by their temperature treatment and sampling period. The dots are surrounded by color corresponding ellipses, which all overlap. Graph B is a boxplot showing the alpha diversity of samples within each temperature treatment and sampling period, where the y-axis is Shannon diversity and the x-axis is the treatment group. The alpha diversity across all groups is similar, with the median of each boxplot between 5 and 5.5.

**Table 1:**
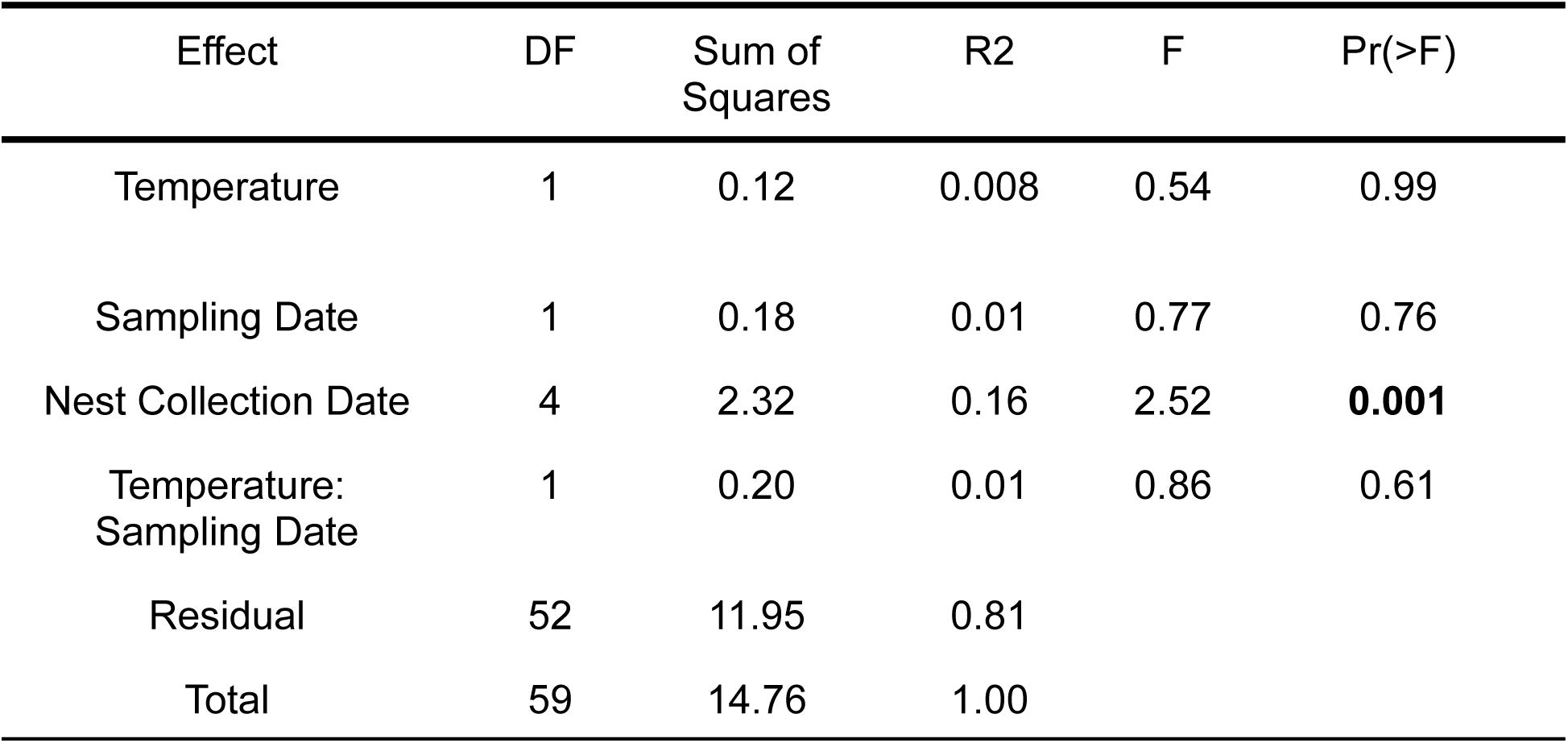
Results of PERMANOVA examining the impact of short-term heatwaves on bacterial communities within pollen provisions, using a Bray-Curtis Distance Matrix.

| Effect | DF | Sum of Squares | R2 | F | Pr(>F) |
| --- | --- | --- | --- | --- | --- |
| Temperature | 1 | 0.12 | 0.008 | 0.54 | 0.99 |
| Sampling Date | 1 | 0.18 | 0.01 | 0.77 | 0.76 |
| Nest Collection Date | 4 | 2.32 | 0.16 | 2.52 | **0.001** |
| Temperature:<br>Sampling Date | 1 | 0.20 | 0.01 | 0.86 | 0.61 |
| Residual | 52 | 11.95 | 0.81 |  |  |
| Total | 59 | 14.76 | 1.00 |  |  |

### Bacterial abundance within pollen provision was consistent across treatments

Neither heat treatment, sampling day, the interaction between treatment and sampling day, nor collection date affected log bacterial copy number per mg pollen provision (all p > 0.3) (Fig. 4).

**Figure 4:**
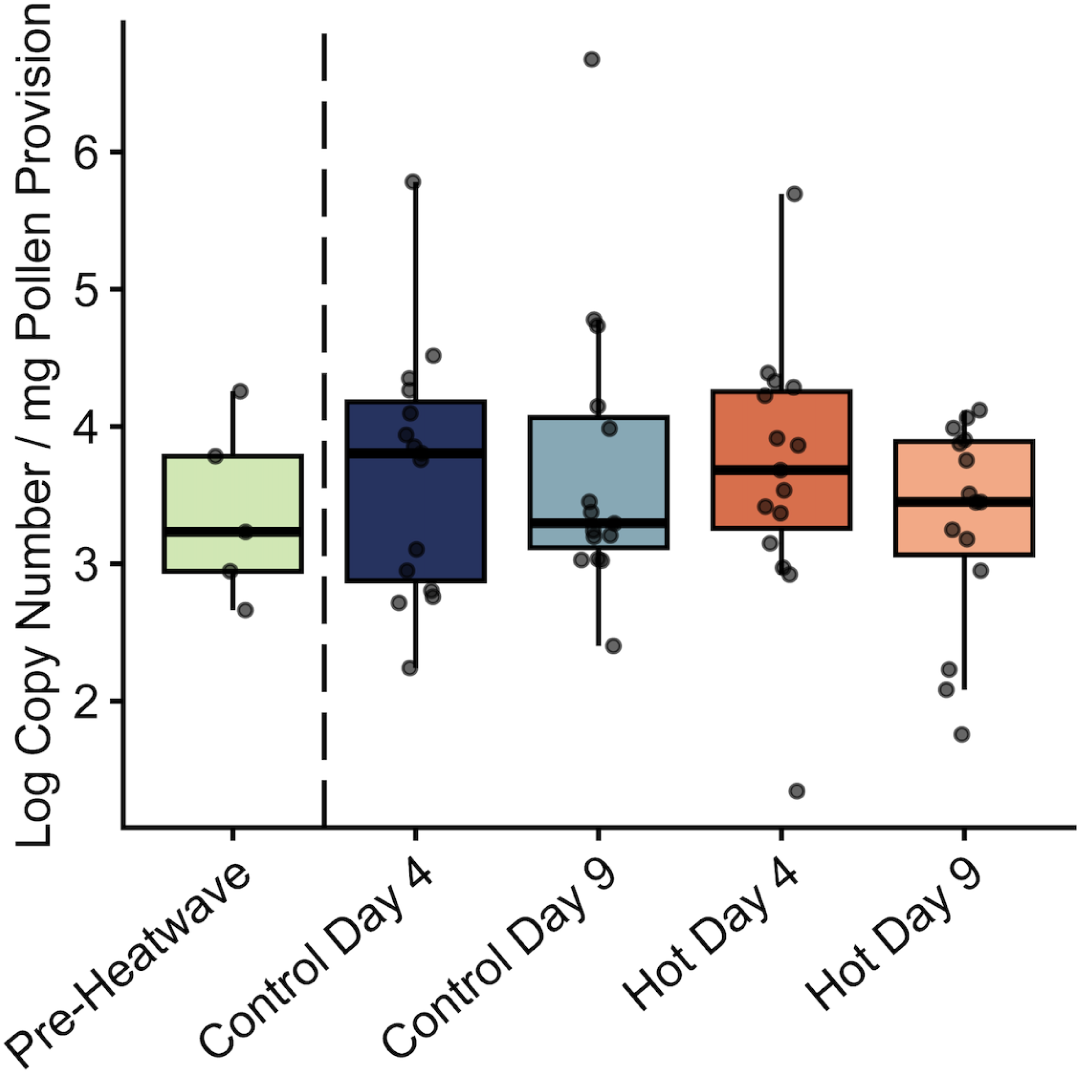
Log bacterial copy number per mg of pollen provision, colored by temperature treatment and sampling day. Pre-heatwave samples are shown here to show the bacterial abundance in the initial community, though we did not include them in the analysis. <u>Alt Text:</u> Boxplots depicting the bacterial abundance within samples, as measured by qPCR. The y-axis is the log copy number per milligram of pollen provision and the x-axis is the temperature treatment – sampling day group (Pre-heatwave, Control Day 4, Control Day 9, Hot Day 4, Hot Day 9). The abundance of bacteria across groups is similar, with the median of each box plot between 3 and 4.

### Larval development was accelerated by heatwave, but mortality was unaffected

The number of bees within each stage was not significantly different at the start of the experiment; however, the proportion of bees in each stage differed across the control and heatwave treatment at Day 4 (χ² = 21.8, df = 3, p < 0.001) and Day 9 (χ² = 11.3, df = 2, p = 0.004), with more bees reaching later stages of larval development in the heat group (Fig. 5). Survival was not impacted by heat treatment, sampling day, or the interaction (all p = 1), with 5/38 offspring dying following heat stress and 2/38 bees dying in the control treatment. All bees died at the egg or first-instar stage, suggesting developmental failure or possible damage during the set-up of treatments.

**Figure 5:**
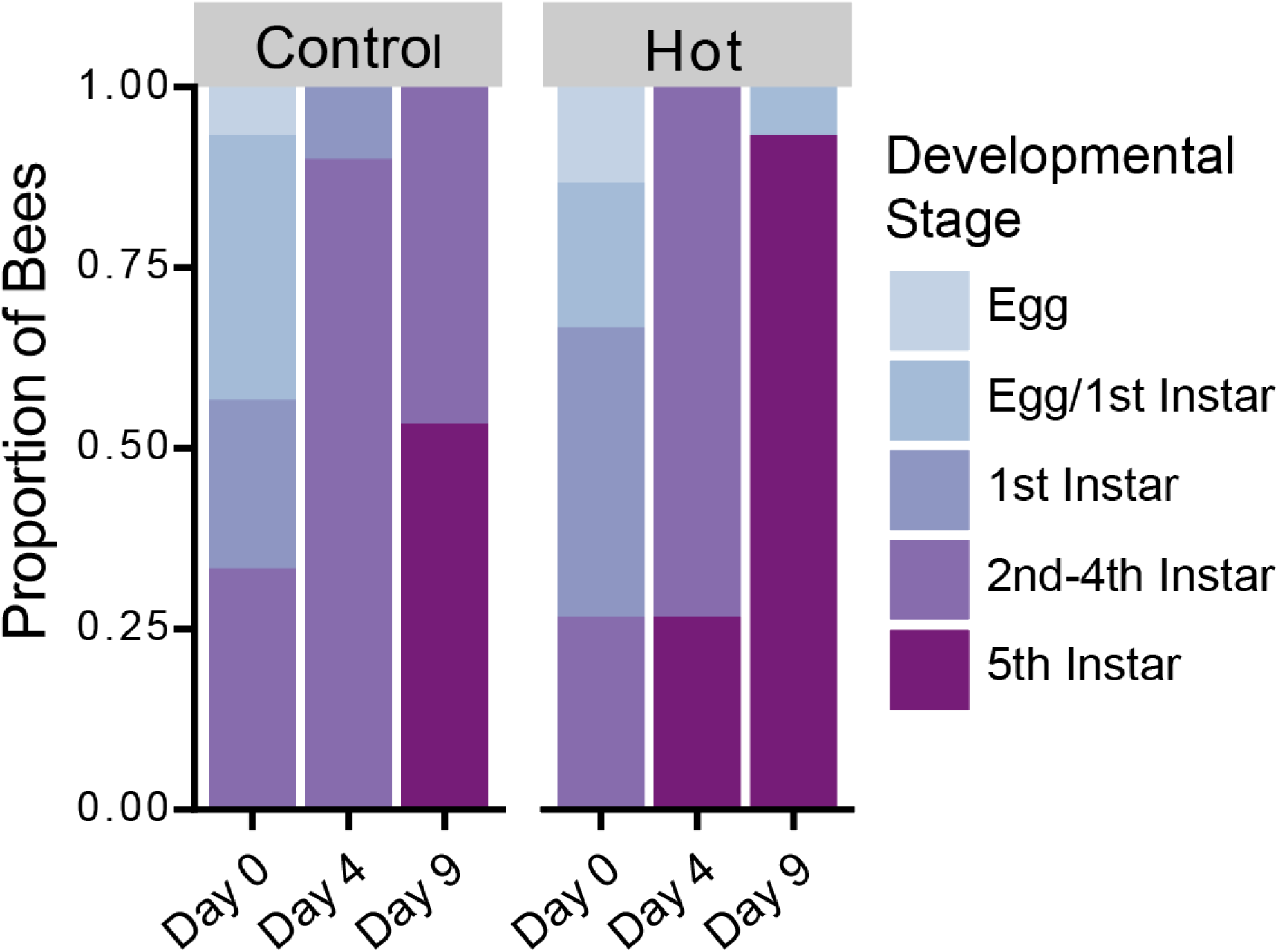
The proportion of blue orchard bees at each developmental stage across the three sampling days, separated by temperature treatment. Day 0 is pre-heatwave, Day 4 is post-heatwave, and Day 9 is post-recovery period. <u>Alt Text:</u> Stacked bar plots showing the proportion of bees in each developmental stage at each sampling day (Day 0, Day 4, Day 9). The y-axis is the proportion of bees in that developmental stage, the x-axis is the sampling day, and the plots are split by temperature treatment (Control or Hot). Though both temperature treatments have similar numbers of bees in early developmental stages, the hot treatment shows increased proportions of bees in the fifth instar at Day 4 and Day 9.

## Discussion

We found that bacterial community composition, diversity, and abundance within blue orchard bee provisions remained unaffected by simulated heatwaves within the range of realistic California temperatures that have previously been shown to impact floral communities (Cecala et al. 2025) and larval health (Melone et al. 2024). Our results differ from previously documented effects of heat on bacteria in flowers (Russell and McFrederick 2022, Cecala et al. 2025) and soil (Bérard et al. 2011, Rocca et al. 2019, Bei et al. 2023). However, like a previous study investigating pollen provision communities subjected to a lower temperature disturbance press (Crowley and Schaeffer 2024), we also failed to detect effects of a heatwave on bacterial abundance or composition. Together, these results suggest that bacterial communities in pollen provisions are resistant to temperature disturbance at the range tested.

One possible hypothesis for temperature resistance is that the dominance of generalist microbial taxa within pollen provisions drives community composition and abundance stability. The bacteria we recovered in blue orchard bee pollen provisions do not seem to be specialized within this system. Close relatives are detected in a variety of environmental sources (e.g., soil, water, sediment, stone) and they do not reach high abundance in resource-rich pollen provisions. Together these factors suggest that they are likely transient, environmentally-derived taxa (Vannette et al. 2025, Martin et al. 2026). Such bacteria may be adapted to survive variable environmental conditions. Indeed, a review of 378 microbial community studies across a multitude of environments (e.g., soil, freshwater, sediment) found that 18% documented robustness to environmental disturbance, and such robustness may be underreported due to biases against publishing null results (Shade et al. 2012). In contrast, specialized microbes may fare worse in the face of climate change due to either increased sensitivity to disturbance or being outcompeted by more resilient generalists (Marvier et al. 2004, Devictor et al. 2008, Clavel et al. 2011, Shan et al. 2017, Renoz et al. 2019, Chen et al. 2021). If such specialist sensitivity is true for provision-associated microbes, then we may expect solitary bees with more host-specialized bacterial associates (e.g., Hammer et al. 2023, Christensen et al. 2024) to be more strongly impacted by climate change stressors (Renoz et al. 2019). Additionally, because the pollen provision microbiome within blue orchard bees is variable across environments and floral sources (Vannette et al. 2025), it is also important to consider how bees from different regions and/or with pollen provisions dominated by floral- or bee-associated taxa (i.e. *Lactobacillus*) may be differentially impacted by disturbance.

Nest collection date was the only factor that impacted bacterial communities in pollen provisions, which may indicate that other environmental components may more strongly shape community composition and diversity. Variation in abiotic factors across dates could filter which microbes are able to survive within an environment. For instance, daily humidity varied across the logged period (May 8, 2025 to May 14, 2025) with some days exhibiting humidity maxima below 80% and others above 90%, which could favor differential survival or growth among bacterial taxa. Additionally, biotic interactions within source pools, such as microbial dispersal or priority effects, could also shape community assembly and maintenance (Vellend 2010, Nemergut et al. 2013). Interestingly, we did not detect any differentially abundant taxa across nest collection dates, which could be due to the fact that the analysis used is unable to account for the variation contributed by temperature treatments or sampling days. These effects could be partitioned by analyzing only pre-heatwave samples; however, we did not sequence enough pre-heatwave samples (n=5) to meet necessary replication. Future studies could more broadly investigate how biotic (i.e. community complexity, priority effects) and abiotic (i.e., temperature, humidity, nutrient availability) environmental factors shape bacterial communities within presumed source pools and subsequent pollen provisions across time when floral resources are held constant.

Bacterial communities within blue orchard bee-pollen provisions were extremely stable over time. Our finding contrasts with a previous study on the temporal dynamics of bacteria within pollen provisions of the horn-face bee (*Osmia cornifrons*), which observed differences in community composition between the initial sampling point and day 9 of the experiment (Kueneman et al. 2023). This difference may be attributed to the communities found within the sampled horn-face bee provisions, which were dominated by fewer, more bee- and plant-specific bacterial taxa (e.g., *Ralstonia*, *Erwinia*, *Sodalis*), whereas blue orchard bee nests sampled here lacked these taxa. It is possible that the high diversity of bacterial communities within our study leads to increased stability over time (the diversity-stability hypothesis), as has been previously proposed in ecological theory (e.g., MacArthur 1955, Lehman et al. 2000) and reported in other systems (e.g., Tilman and Downing 1994, Tilman 1996, Rodrigues et al. 2025).

Although many examples of the diversity-stability hypothesis are related to functional redundancy within the community (i.e. multiple species perform the same function, enabling overall community stability even if a member is lost), it is also possible that high diversity promotes competition between species thereby preventing a single species from dominating the community. Bacterial diversity may also be important for host health, as more diverse bacterial communities in pollen provisions are positively correlated with blue orchard bee developmental success (Westreich et al. 2023). Alternatively, it is possible that bacterial diversity, and thus positive host health implications, reflect broader trends of resource availability or diversity (Vannette et al. 2025).

Although the bacterial communities of provisions were not affected by temperature stress and the heatwave was not extreme enough to result in larval death, bee larvae experienced marked changes in development. Larvae increased the speed of development when subjected to a 35 °C heatwave, resulting in more larvae in the fifth instar immediately after the temperature stress. Even following the return of heat-exposed bees to control temperatures for 5 days, these larvae continued to develop more rapidly than bees kept under constant control temperatures for the entire experiment. These data compliment trends from Melone et al. 2024, in which blue orchard bee larvae subjected to a 31 °C heatwave trended towards faster development than those kept at a constant 25 °C, while larvae exposed to a 37 °C heatwave took significantly longer to develop and experienced high mortality. Together, this suggests that heatwaves accelerate development up until an optimal temperature at which point heat stress begins to significantly impair development speed; however, it is important to note that we did not follow larvae to cocoon completion.

In conclusion, we found that bacterial communities within blue orchard bee pollen provisions are not affected by short-term heatwaves despite clear developmental impacts on larvae. In combination with previous literature (Crowley and Schaeffer 2024), this suggests that bacteria within provisions do not enhance or reduce blue orchard bee survival to heatwaves. Although these bacteria must exhibit a thermal maxima, it appears to be outside the temperature range of blue orchard bee larvae. As a result, heat stress is likely to result in larval death before relationships between blue orchard bees and the bacteria within their pollen provision break down. Instead, the impacts of disturbance on bee fitness could be mediated by relationships with fungi as seen in other bee-disturbance contexts (reviewed in Rutkowski et al. 2023), or endogenous to bee physiology. Across bee species more broadly, sensitivity to heat varies based on numerous factors, such as nesting behavior (Da Silva et al. 2026), sociality (Vilchez-Russell and Rafferty 2024), urbanization, and pesticide exposure, though research on bee thermal tolerance remains limited (reviewed in Baena-Díaz et al. 2026). Because heat stress can disrupt mutualistic and facultative relationships between other insect-bacteria partnerships (Dunbar et al. 2007, Burke et al. 2010, Kikuchi et al. 2016, Ross et al. 2017, Shan et al. 2017, Renoz et al. 2019) and different microbial taxa (i.e. fungi, viruses) may be differentially impacted by heat stress (Philippot et al. 2021), it remains critical to understand how life history traits of different bee species impact their relationships with diverse microbial partners under disturbance. Overall, our study finds that blue orchard bees are sensitive to heatwaves, but this effect is not exacerbated by a bacterial Achilles’ Heel.

## Supporting information

supplemental methods and results

## Acknowledgements

The authors would like to thank members of the Vannette Lab for feedback on the manuscript, as well as Helen Noroian, Alex Ma, Campbell Sinclair, Joseph Tauser, and the Integrated Microbiome Resource at Dalhousie University for technical project support.

## Data Availability

The sequence data used in this paper was submitted to the NCBI Sequence Reads Archive (SRA) under BioProject ID PRJNA1498200. The code used to process all data and all data files can be found on the lead author’s github at the following URL: https://github.com/lexienichole/OsmiaGlobalChange2026. All files will be made publicly available upon publication.

## References

Altschul SF, Gish W, Miller W, et al. 1990. Basic local alignment search tool. J. Mol. Biol. 215(3):403–410. 10.1016/S0022-2836(05)80360-2

Anguita-Maeso M, Haro C, Navas-Cortés JA, et al. 2022. Primer Choice and Xylem-Microbiome-Extraction Method Are Important Determinants in Assessing Xylem Bacterial Community in Olive Trees. Plants. 11(10):1320. 10.3390/plants11101320

Baena-Díaz F, Morales-Gómez M, Ratoni B, et al. 2026. General patterns and predictors of thermal tolerance in bees: A global synthesis across taxa and environments. J. Therm. Biol. 140:104538. 10.1016/j.jtherbio.2026.104538

Bei Q, Reitz T, Schnabel B, et al. 2023. Extreme summers impact cropland and grassland soil microbiomes. ISME J. 17(10):1589–1600. 10.1038/s41396-023-01470-5

Bérard A, Bouchet T, Sévenier G, et al. 2011. Resilience of soil microbial communities impacted by severe drought and high temperature in the context of Mediterranean heat waves. Eur. J. Soil Biol. 47(6):333–342. 10.1016/j.ejsobi.2011.08.004

Boyle JA, Murphy BK, Ensminger I, et al. 2024. Resistance and resilience of soil microbiomes under climate change. Ecosphere. 15(12):e70077. 10.1002/ecs2.70077

Brar G, Floden M, McFrederick Q, et al. 2024. Environmentally acquired gut-associated bacteria are not critical for growth and survival in a solitary bee, Megachile rotundata. Appl. Environ. Microbiol. 90(9):e02076–23. 10.1128/aem.02076-23

Burke G, Fiehn O, Moran N. 2010. Effects of facultative symbionts and heat stress on the metabolome of pea aphids. ISME J. 4(2):242–252. 10.1038/ismej.2009.114

Callahan B. 2024. Silva taxonomic training data formatted for DADA2 (Silva version 138.2) [Data set]. Zenodo. 10.5281/zenodo.14169026

Callahan BJ, McMurdie PJ, Rosen MJ, et al. 2016. DADA2: High-resolution sample inference from Illumina amplicon data. Nat. Methods. 13(7):581–583. 10.1038/nmeth.3869

CaraDonna PJ, Cunningham JL, Iler AM. 2018. Experimental warming in the field delays phenology and reduces body mass, fat content and survival: Implications for the persistence of a pollinator under climate change. Funct. Ecol. 32(10):2345–2356. 10.1111/1365-2435.13151

Cardoso P, Barton PS, Birkhofer K, et al. 2020. Scientists’ warning to humanity on insect extinctions. Biol. Conserv. 242:108426. 10.1016/j.biocon.2020.108426

Cecala JM, Landucci L, Vannette RL. 2025. Seasonal Assembly of Nectar Microbial Communities Across Angiosperm Plant Species: Assessing Contributions of Climate and Plant Traits. Ecol. Lett. 28(1):e70045. 10.1111/ele.70045

Chelius MK, Triplett EW. 2001. The Diversity of Archaea and Bacteria in Association with the Roots of Zea mays L. Microb. Ecol. 41(3):252–263. 10.1007/s002480000087

Chen Y-J, Leung PM, Wood JL, et al. 2021. Metabolic flexibility allows bacterial habitat generalists to become dominant in a frequently disturbed ecosystem. ISME J. 15(10):2986–3004. 10.1038/s41396-021-00988-w

Christensen SM, Srinivas SN, McFrederick QS, et al. 2024. Symbiotic bacteria and fungi proliferate in diapause and may enhance overwintering survival in a solitary bee. ISME J. 18(1):wrae089. 10.1093/ismejo/wrae089

Clavel J, Julliard R, Devictor V. 2011. Worldwide decline of specialist species: toward a global functional homogenization? Front. Ecol. Environ. 9(4):222–228. 10.1890/080216

Colwell RK, Guzman LM, Steinke D, et al. 2026. Constructing a lower-bound estimate of the global number of insect species on a hyperdiverse empirical foundation. Proc. Natl. Acad. Sci. 123(27):e2524283123. 10.1073/pnas.2524283123

Cornwallis CK, van ‘t Padje A, Ellers J, et al. 2023. Symbioses shape feeding niches and diversification across insects. Nat. Ecol. Evol. 7(7):1022–1044. 10.1038/s41559-023-02058-0

Created in BioRender. Martin, L. (2027) https://BioRender.com/rdmrsdj

Crowley BL, Schaeffer RN. 2024. Bee microbiomes in a changing climate: Investigating the effects of temperature on solitary bee life history and health. Environ. Microbiol. 26(11):e70002. 10.1111/1462-2920.70002

Da Silva CRB, Beaman JE, Dorey JB, et al. 2026. Nesting behaviour predicts heat tolerance evolution and climate vulnerability in bees. Nat. Commun. 17(1):4961. 10.1038/s41467-026-73689-7

Davis NM, Proctor DM, Holmes SP, et al. 2018. Simple statistical identification and removal of contaminant sequences in marker-gene and metagenomics data. Microbiome. 6(1):226. 10.1186/s40168-018-0605-2

Devictor V, Julliard R, Jiguet F. 2008. Distribution of specialist and generalist species along spatial gradients of habitat disturbance and fragmentation. Oikos. 117(4):507–514. 10.1111/j.0030-1299.2008.16215.x

Dharampal PS, Carlson C, Currie CR, et al. 2019. Pollen-borne microbes shape bee fitness. Proc. R. Soc. B Biol. Sci. 286(1904):20182894. 10.1098/rspb.2018.2894

Dharampal PS, Danforth BN, Steffan SA. 2022. Exosymbiotic microbes within fermented pollen provisions are as important for the development of solitary bees as the pollen itself. Ecol. Evol. 12(4). 10.1002/ece3.8788

Dharampal PS, Hetherington MC, Steffan SA. 2020. Microbes make the meal: oligolectic bees require microbes within their host pollen to thrive. Ecol. Entomol. 45(6):1418–1427. 10.1111/een.12926

Dunbar HE, Wilson ACC, Ferguson NR, et al. 2007. Aphid Thermal Tolerance Is Governed by a Point Mutation in Bacterial Symbionts. PLOS Biol. 5(5):e96. 10.1371/journal.pbio.0050096

Engl T, Eberl N, Gorse C, et al. 2018. Ancient symbiosis confers desiccation resistance to stored grain pest beetles. Mol. Ecol. 27(8):2095–2108. 10.1111/mec.14418

Gentleman R, Carey V, Huber W, et al. 2024. genefilter: methods for filtering genes from high-throughput experiments. R package version 1.88.0. 10.18129/B9.bioc.genefilter

Gruntenko NE, Ilinsky YYu, Adonyeva NV, et al. 2017. Various Wolbachia genotypes differently influence host Drosophila dopamine metabolism and survival under heat stress conditions. BMC Evol. Biol. 17. 10.1186/s12862-017-1104-y

Gudowska A, Tofilski A, Moroń D. 2026. Thermal carry-over: lasting impacts of early life heat stress on morphology and flight performance in solitary bee Osmia bicornis. Proc. R. Soc. B Biol. Sci. 293(2074):20260069. 10.1098/rspb.2026.0069

Hammer TJ, Kueneman J, Argueta-Guzmán M, et al. 2023. Bee breweries: The unusually fermentative, lactobacilli-dominated brood cell microbiomes of cellophane bees. Front. Microbiol. 14:1114849. 10.3389/fmicb.2023.1114849

Kazenel MR, Wright KW, Griswold T, et al. 2024. Heat and desiccation tolerances predict bee abundance under climate change. Nature. 628(8007):342–348. 10.1038/s41586-024-07241-2

Keller A, Grimmer G, Steffan-Dewenter I. 2013. Diverse Microbiota Identified in Whole Intact Nest Chambers of the Red Mason Bee Osmia bicornis (Linnaeus 1758). PLoS ONE. 8(10). 10.1371/journal.pone.0078296

Kikuchi Y, Tada A, Musolin DL, et al. 2016. Collapse of Insect Gut Symbiosis under Simulated Climate Change. mBio. 7(5):10.1128/mbio.01578-16. 10.1128/mbio.01578-16

Kueneman JG, Gillung J, Van Dyke MT, et al. 2023. Solitary bee larvae modify bacterial diversity of pollen provisions in the stem-nesting bee, Osmia cornifrons (Megachilidae). Front. Microbiol. 13. 10.3389/fmicb.2022.1057626

Kuhlmann M, Guo D, Veldtman R, et al. 2012. Consequences of warming up a hotspot: species range shifts within a centre of bee diversity. Divers. Distrib. 18(9):885–897. 10.1111/j.1472-4642.2011.00877.x

LeBuhn G, Vargas Luna J. 2021. Pollinator decline: what do we know about the drivers of solitary bee declines? Curr. Opin. Insect Sci. 46:106–111. 10.1016/j.cois.2021.05.004

Lemoine MM, Engl T, Kaltenpoth M. 2020. Microbial symbionts expanding or constraining abiotic niche space in insects. Curr. Opin. Insect Sci. 39:14–20. 10.1016/j.cois.2020.01.003

Love M, Huber W, Anders S. 2014. Moderated estimation of fold change and dispersion for RNA-seq data with DESeq2. Genome Biology. 12(15):550.

Martin AN, Stuligross C, Williams NM, et al. 2026. Floral microbes provisioned by Osmia lignaria establish in larval food stores, but do not affect bee development or survival. FEMS Microbiol. Ecol. 102(4):fiag025. 10.1093/femsec/fiag025

Marvier M, Kareiva P, Neubert MG. 2004. Habitat Destruction, Fragmentation, and Disturbance Promote Invasion by Habitat Generalists in a Multispecies Metapopulation. Risk Anal. 24(4):869–878. 10.1111/j.0272-4332.2004.00485.x

McMurdie PJ, Holmes S. 2013. phyloseq: An R Package for Reproducible Interactive Analysis and Graphics of Microbiome Census Data. PLOS ONE. 8(4):e61217. 10.1371/journal.pone.0061217

Meehl GA, Arblaster JM, Tebaldi C. 2007. Contributions of natural and anthropogenic forcing to changes in temperature extremes over the United States. Geophys. Res. Lett. 34(19). 10.1029/2007GL030948

Melone GG, Stuligross C, Williams NM. 2024. Heatwaves increase larval mortality and delay development of a solitary bee. Ecol. Entomol. 49(3):433–444. 10.1111/een.13317

Nemergut DR, Schmidt SK, Fukami T, et al. 2013. Patterns and Processes of Microbial Community Assembly. Microbiol. Mol. Biol. Rev. 77(3):342–356. 10.1128/mmbr.00051-12

Nguyen PN, Rehan SM. 2022. Developmental microbiome of the small carpenter bee, Ceratina calcarata. Environ. DNA. 4(4):808–819. 10.1002/edn3.291

Philippot L, Griffiths BS, Langenheder S. 2021. Microbial Community Resilience across Ecosystems and Multiple Disturbances. Microbiol. Mol. Biol. Rev. 85(2):e00026–20. 10.1128/MMBR.00026-20

Potts SG, Biesmeijer JC, Kremen C, et al. 2010. Global pollinator declines: trends, impacts and drivers. Trends Ecol. Evol. 25(6):345–353. 10.1016/j.tree.2010.01.007

Quast C, Pruesse E, Yilmaz P, et al. 2013. The SILVA ribosomal RNA gene database project: improved data processing and web-based tools. Nucleic Acids Res. 41(D1):D590–D596. 10.1093/nar/gks1219

R Core Team. 2024. R: A language and environment for statistical computing. R Foundation for Statistical Computing, Vienna, Austria. http://www.R-project.org/. [accessed 2025 July 17]. https://cir.nii.ac.jp/crid/1574231874043578752.

Renoz F, Pons I, Hance T. 2019. Evolutionary responses of mutualistic insect–bacterial symbioses in a world of fluctuating temperatures. Curr. Opin. Insect Sci. 35:20–26. 10.1016/j.cois.2019.06.006

Rocca JD, Simonin M, Blaszczak JR, et al. 2019. The Microbiome Stress Project: Toward a Global Meta-Analysis of Environmental Stressors and Their Effects on Microbial Communities. Front. Microbiol. 9. 10.3389/fmicb.2018.03272

Rodrigues AV, Rissanen T, Jones MM, et al. 2025. Cross-Taxa Analysis of Long-Term Data Reveals a Positive Biodiversity-Stability Relationship With Taxon-Specific Mechanistic Underpinning. Ecol. Lett. 28(4):e70003. 10.1111/ele.70003

Ross PA, Wiwatanaratanabutr I, Axford JK, et al. 2017. Wolbachia Infections in Aedes aegypti Differ Markedly in Their Response to Cyclical Heat Stress. PLOS Pathog. 13(1):e1006006. 10.1371/journal.ppat.1006006

Rothman JA, Andrikopoulos C, Cox-Foster D, et al. 2019. Floral and Foliar Source Affect the Bee Nest Microbial Community. Microb. Ecol. 78(2):506–516. 10.1007/s00248-018-1300-3

Rothman JA, Leger L, Graystock P, et al. 2019. The bumble bee microbiome increases survival of bees exposed to selenate toxicity. Environ. Microbiol. 21(9):3417–3429. 10.1111/1462-2920.14641

Russell KA, McFrederick QS. 2022. Floral nectar microbial communities exhibit seasonal shifts associated with extreme heat: Potential implications for climate change and plant-pollinator interactions. Front. Microbiol. 13. 10.3389/fmicb.2022.931291

Rutkowski D, Weston M, Vannette RL. 2023. Bees just wanna have fungi: a review of bee associations with nonpathogenic fungi. FEMS Microbiol. Ecol. 99(8):1–16.

Shade A, Peter H, Allison SD, et al. 2012. Fundamentals of Microbial Community Resistance and Resilience. Front. Microbiol. 3. 10.3389/fmicb.2012.00417

Shan H-W, Deng W-H, Luan J-B, et al. 2017. Thermal sensitivity of bacteriocytes constrains the persistence of intracellular bacteria in whitefly symbiosis under heat stress. Environ. Microbiol. Rep. 9(6):706–716. 10.1111/1758-2229.12554

[Software] Oksanen J, Simpson GL, Blanchet FG, et al. 2024. vegan: Community Ecology Package. [accessed 2024 Dec 13]. https://cran.r-project.org/web/packages/vegan/index.html.

Soroye P, Newbold T, Kerr J. 2020. Climate change contributes to widespread declines among bumble bees across continents. Science. 367(6478):685–688. 10.1126/science.aax8591

Tilman D. 1996. Biodiversity: Population Versus Ecosystem Stability. Ecology. 77(2):350–363. 10.2307/2265614

Tilman D, Downing JA. 1994. Biodiversity and stability in grasslands. Nature. 367(6461):363–365. 10.1038/367363a0

Torchio PF. 1989. In-Nest Biologies and Development of Immature Stages of Three Osmia Species (Hymenoptera: Megachilidae). Ann. Entomol. Soc. Am. 82(5):599–615.

Ummenhofer CC, Meehl GA. 2017. Extreme weather and climate events with ecological relevance: a review. Philos. Trans. R. Soc. B Biol. Sci. 372(1723):20160135. 10.1098/rstb.2016.0135

Vannette RL, Williams NM, Peterson SS, et al. 2025. Pollen diet, more than geographic distance, shapes provision microbiome composition in two species of cavity-nesting bees. FEMS Microbiol. Ecol. 101(8):fiaf067. 10.1093/femsec/fiaf067

Vellend M. 2010. Conceptual Synthesis in Community Ecology. Q. Rev. Biol. 85(2):183–206. 10.1086/652373

Vilchez-Russell KA, Rafferty NE. 2024. Effects of heat shocks, heat waves, and sustained warming on solitary bees. Front. Bee Sci. 2. 10.3389/frbee.2024.1392848

Voulgari-Kokota A, McFrederick QS, Steffan-Dewenter I, et al. 2019. Drivers, Diversity, and Functions of the Solitary-Bee Microbiota. Trends Microbiol. 27(12):1034–1044. 10.1016/j.tim.2019.07.011

Voulgari-Kokota A, Steffan-Dewenter I, Keller A. 2020. Susceptibility of Red Mason Bee Larvae to Bacterial Threats Due to Microbiome Exchange with Imported Pollen Provisions. Insects. 11(6):373. 10.3390/insects11060373

Westreich LR, Westreich ST, Tobin PC. 2023. Bacterial and Fungal Symbionts in Pollen Provisions of a Native Solitary Bee in Urban and Rural Environments. Microb. Ecol. 86(2):1416–1427. 10.1007/s00248-022-02164-9

