## Supplementary material for "Heatwaves do not impact bacteria within pollen provisions, despite accelerating blue orchard bee (*Osmia lignaria*) larval development": Fig S.

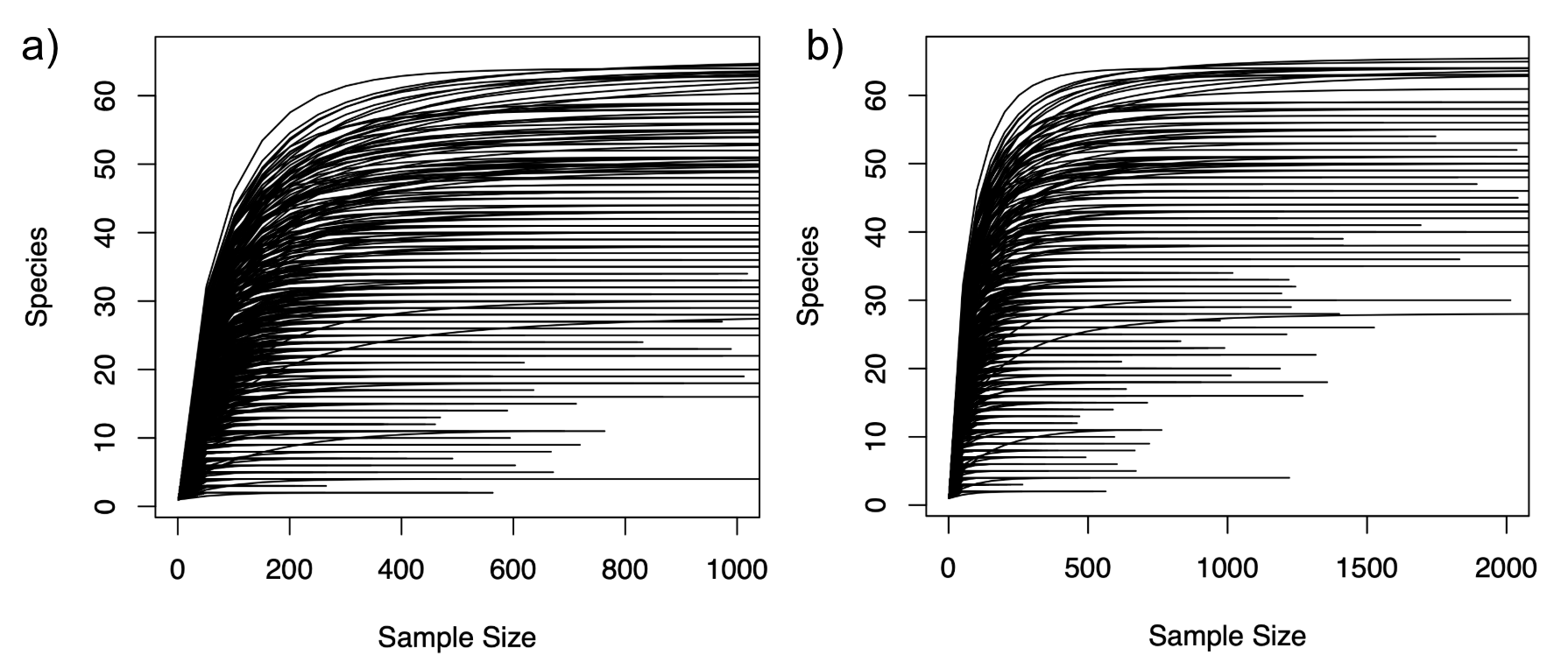

**Figure S1**: Sampling curves for the bacterial dataset with legend restricted to (a) 1000 reads and (b) 2000 reads. Curves were used to determine that samples had sufficient sampling depth.

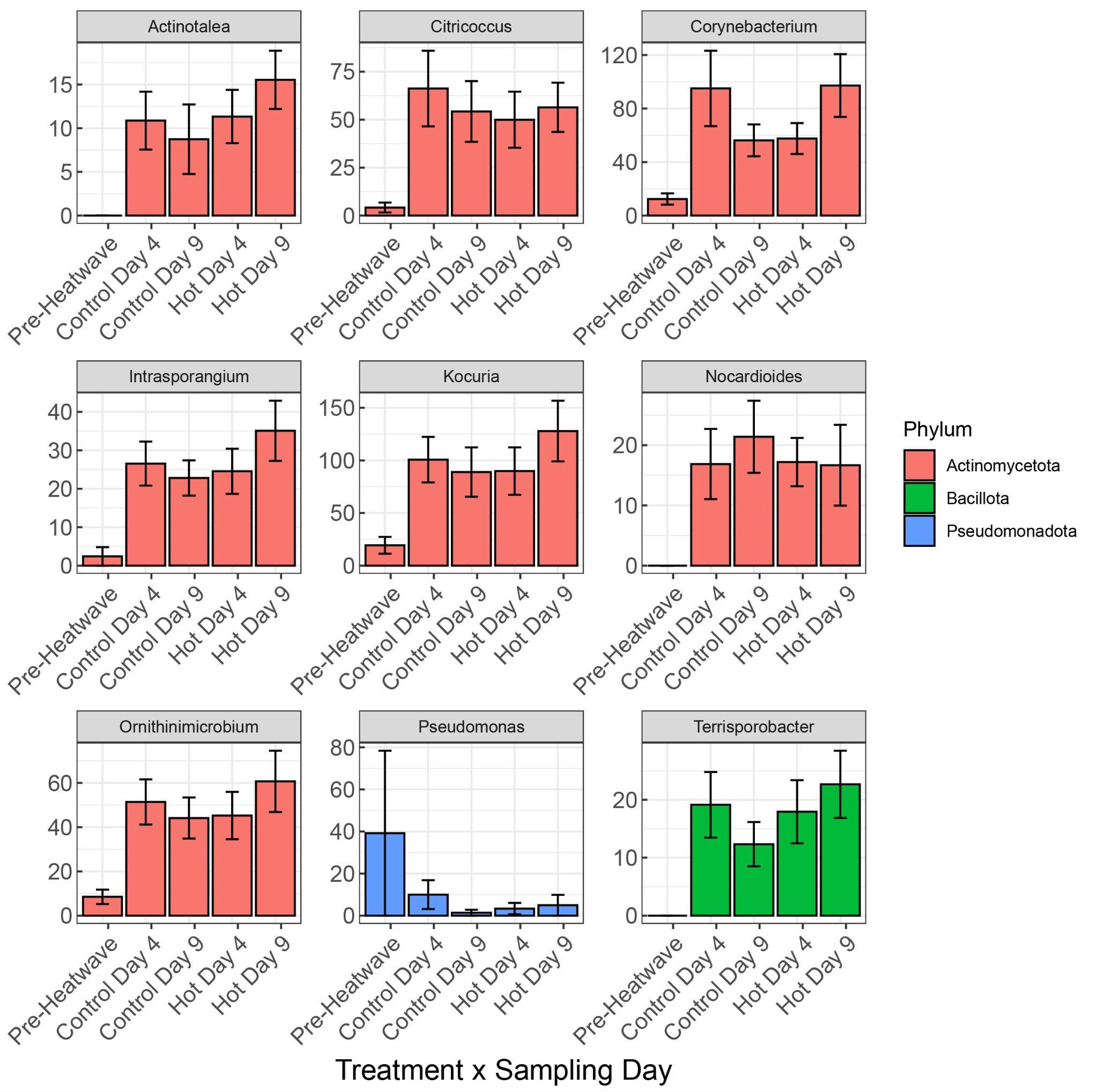

**Figure S2**: DESeq plot highlighting the bacterial genera identified as having differential relative abundance across temperature treatment-sampling day categories.
