## supplemental methods and results for "Heatwaves do not impact bacteria within pollen provisions, despite accelerating blue orchard bee (*Osmia lignaria*) larval development"

Analyzing bacterial communities within pollen provisions over time

*Statistical Analysis*

To investigate if overall bacterial communities differed from the initial bacterial community over time, we ran a PERMANOVA using the adonis2() function in R (“vegan” package; (Oksanen et al. 2024) with combined temperature treatment-sampling period categories (Day 0, Heat Treatment Day 4, Heat Treatment Day 9, Control Treatment Day 4, Control Treatment Day 9) and nest collection date as predictors. To investigate if alpha diversity changed over time across treatment, we performed a linear model with the same variables described above as predictors and Shannon’s diversity as the response variable. To determine if the abundance of bacterial communities in pollen provision shifts over time, we ran a linear mixed model with log copy number per mg as the response variable, the same predictors described above main effects, and qPCR plate ID as a random effect.

*Results*

There were no differences in bacterial community composition, alpha diversity, or bacterial abundance across the combined treatment-sampling period categories, suggesting that these factors did not vary over time. The nest collection date significantly impacted bacterial community composition over time (F_5,55_  = 2.45, p = 0.001), though dispersion differed across dates (betadisper F = 11.3, df = 5, P = 0.001). There was no impact of nest collection on bacterial alpha diversity or abundance within pollen provisions.
